# Differential recognition and cytokine induction by the peptidorhamnomannan from *Sporothrix brasiliensis* and *S. schenckii*

**DOI:** 10.1101/2022.01.09.475552

**Authors:** Brenda Kischkel, Leila Lopes-Bezerra, Carlos P. Taborda, Leo A. B. Joosten, Jéssica Cristina dos Santos, Mihai G. Netea

**Author notes:** Lead contact, These authors jointly supervised this work: Jéssica Cristina dos Santos and Mihai G. Netea. Corresponding authors: Mihai G. Netea, M.D PhD and Jéssica Cristina dos Santos, PhD, Department of Internal Medicine, Radboud University Medical Center, 6500 HB Nijmegen, the Netherlands, and.

## Abstract

Sporotrichosis is a deep mycosis caused by dimorphic species of the genus *Sporothrix*, with differences in pathogenicity between *S. schenckii* and *S. brasiliensis* species. Recently, it was discovered that the cell wall peptidorhamnomannan (PRM) of *Sporothrix spp*. is a pathogen associated molecular pattern (PAMP). Interestingly, *S. brasiliensis* PRM has additional unknown rhamnose residues. We hypothesize that the structural differences of *Sporothrix* spp PRMs impact the host’s immune response and may explain the severity of sporotrichosis caused by *S. brasiliensis*. Here we demonstrate that *S. brasiliensis* yeasts and its PRM (*S.b* PRM) induced a strong inflammatory response in human PBMCs, with high production of TNF-α, IL-6 and IL-1β and induction of T-helper cytokines IFN-γ, IL-17 and IL-22. In contrast, *S. schenckii* yeasts and its PRM induced higher concentrations of interleukin-1 receptor antagonist (IL-1Ra), which resulted in low production of T-helper cytokines such as IL-17 and IL-22. CR3 and dectin-1 were required for cytokine induction by both PRMs, while TLR2 and TLR4 were required for the response of *S.s* PRM and *S.b* PRM, respectively. IL-1β and IL-1α production induced by *S. brasiliensis* yeasts and *S.b* PRM were dependent on inflammasome and caspase-1 activation. *S. schenckii* and *S.s* PRM were able to induce IL-1β independent of ROS. In conclusion, these findings improve our understanding of the pathogenesis of *Sporothrix* spp. by reporting differences of immunological responses induced by *S. schenckii* and *S. brasiliensis*. The study also opens the gateway for novel treatment strategies targeting local inflammation and tissue destruction induced by *S. brasiliensis* infection through IL-1 inhibition.

## Introduction

Sporotrichosis is a deep mycosis caused by the traumatic inoculation into the skin of infectious propagules of pathogenic *Sporothrix* species, *Sporothrix schenckii, Sporothrix globosa* and *Sporothrix brasiliensis*. The classical infection caused by *S. schenckii* affects mucous membranes, lymphatic vessels and osteoarticular sites (Queiroz-Telles et al., 2011). The disease is a granulomatous fungal infection which can be manifested as subcutaneous, extracutaneous and disseminated clinical forms and it is known to cause disease in both immunocompromised and immunocompetent individuals (Lopes-Bezerra et al., 2018a; Queiroz-Telles et al., 2019). *S. schenckii* is a species of worldwide occurrence, popularly known as ‘*rose gardener’s disease*’, since plants were the main source of fungus’ transmission (sapronotic route) (Lopes-Bezerra et al., 2006). For a century, sporotrichosis was attributed to *S. schenckii* as a single etiological agent. However, in 2007, the knowledge about the epidemiology of sporotrichosis has taken an important step forward with the description of cryptic pathogenic species such as *S. brasiliensis and S. globosa* (Marimon et al., 2007) and the zoonotic transmission of *S. brasiliensis* and *S. schenckii* (Gremião et al., 2017; Han and Kano, 2021). In particular, *S. brasiliensis* emerged in Brazil causing severe and lethal infections in cats, that led to massive transmissions to humans (zoonotic route) (Gremião et al., 2020). *S. brasiliensis* infection is currently considered as an hyperendemic zoonosis in Brazil (Gremião et al., 2020). As it continues to expand to other countries in Latin America, *S. brasiliensis* infection has become an urgent public health problem (Etchecopaz et al., 2021; Queiroz-Telles et al., 2019).

Studies comparing the virulence of the two species have shown that *S. brasiliensis* is more virulent than *S. sckenckii* in a murine model of subcutaneous infection. Mice infected with *S. brasiliensis* produced higher amounts of cytokines and developed long-term chronic infection and systemic spread of the fungus in which was associated with severe granulomatous lesions. On the other hand, mice infected with *S. schenckii* quickly controlled the infection better, with a reduced local and systemic fungal load (Batista-Duharte et al., 2018). Of importance, the severe and fatal cases described in human patients were attributed to infection caused by *S. brasiliensis* (Freitas et al., 2015; Orofino-Costa et al., 2013; Silva-Vergara et al., 2012). Currently, the antifungal drugs commonly used to treat sporotrichosis are itraconazole, potassium iodide and amphotericin B. Unfortunately, recent studies demonstrate that *S. brasiliensis* is less susceptible to the azole class of antifungals than *S. schenckii* (Borba-Santos et al., 2015; Fernández-Silva et al., 2014).

The molecular substrate of the differences in virulence and pathogenicity between the main *Sporothrix* species is poorly known, although recent studies have reported differences between the cell wall components of *S. brasiliensis* and *S. schenckii* (Lopes-Bezerra et al., 2018b; Neves et al., 2021). The cell wall of both S*porothrix* species is mainly composed of β-glucans (1–3, 1–6 and 1–4 bonds), chitin and a peptidorhamnomannan (PRM) (Lopes-Bezerra et al., 2018b). Rhamnomannan corresponds to the N-glycans (polysaccharide chain) isolated by hot alkali extraction and has exactly the same structure, Rha*p* α 1-3 Man*p* lateral chains, in both species (Villalobos-Duno et al., 2021). The peptidorhamnomannan is the glycopeptide component of the cell wall and is composed by a peptide core decorated with N-glycans and O-glycans (Lopes-Bezerra, 2011). *S. brasiliensis* has a thicker cell wall, which is 1.5 and 2 fold richer in chitin and rhamnose, respectively, in comparison to *S. schenckii*. Specifically, ^1^H ^13^C NMR (nuclear magnetic resonance) spectra showed unique signals correspondent to rhamnose units in the *S. brasiliensis* PRM structure that are not observed in *S. schenckii* (Neves et al., 2021). Therefore, *S. brasiliensis* PRM has common structures found in *S. schenckii* PRM but also unique rhamnose epitopes. In turn, this may explain the different clinical presentation between infections caused by the two *Sporothrix* species.

In the present study, we investigated the ability of the PRM isolated from the cell wall of *S. schenckii* and *S. brasiliensis*, to induce inflammation, in comparison with the immune response induced by the intact yeasts of both species. We have identified IL-1β as a key cytokine for inducing an inflammatory response caused by *S. brasiliensis* and *S.b* PRM. Moreover, we identified that *S. schenkii* and *S.s* PRM induces higher concentrations of interleukin-1 antagonist receptor (IL-1Ra) than *S. brasiliensis*. Inhibition of IL-1 receptor, inflammasome or caspase-1 were able to decrease the production of proinflammatory cytokines by *S. brasiliensis*. Since one of the main characteristics of the disease is the exacerbation of the inflammatory response that leads to the destruction of the host’s skin, our findings contribute to the understanding of the pathogenesis of *Sporothrix* spp. and offer possibilities for improved treatment of sporotrichosis.

## Results

### Differential induction of inflammatory cytokines by *Sporothrix brasiliensis* and *Sporothrix schenckii*

We evaluated the production of pro- and anti-inflammatory innate immune cytokines in human PBMCs after 24 h exposure with *S. brasiliensis* PRM (*S.b* PRM), *S. schenckii* PRM (*S.s* PRM) as well as heat-killed yeasts of *S. brasiliensis, S. schenckii* and *Candida albicans* as control stimuli. *S. brasiliensis* yeasts induced higher levels of IL-1β, TNF-α and IL-6 than *S. schenckii* and *C. albicans* yeasts (**Figure 1 A, B** and **C**). On the other hand, no differences were observed in TNF-α production induced by *S. schenckii* or *C. albicans* (**Figure 1 B**). In contrast, production of IL-1Ra was significantly higher in cells stimulated with *S. schenckii* than *S. brasiliensis* (**Figure D**).

**Figure 1.**
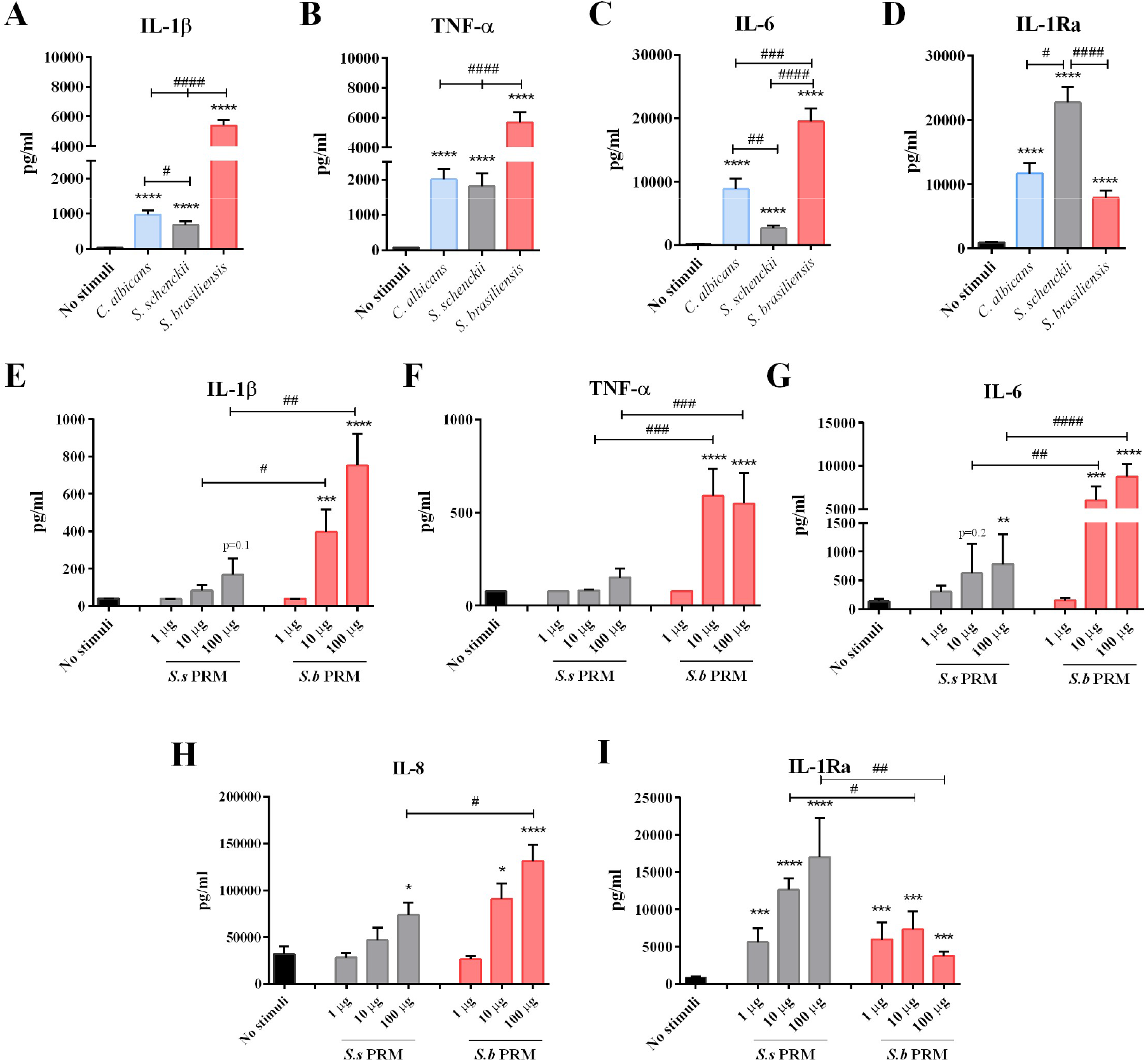
Stimulation of PBMCs with *S. brasiliensis* and *S.b* PRM results in high production of pro-inflammatory cytokines, while *S. schenckii* and S.s PRM induces IL-1Ra. PBMCs were stimulated with heat-killed *Candida albicans, S. schenckii* and *S. brasiliensis*. After 24 hours of stimulation, supernatants were collected and **A.** IL-1β (n=8), **B.** TNF-α (n=8), **C.** IL-6 (n=8), and **D.** IL-1Ra (n=8) production were measured by ELISA. PBMCs were exposed to different concentrations (1, 10 and 100 μg/ml) of peptidorhamnomannans (PRMs) for 24 hours. After 24 hours of stimulation, **E.** IL-1β (n=7) **F.** TNF-α (n=7), **G.** IL-6 (n=7), **H.** IL-8 (n=8), and **I.** IL-1Ra (n=8) production were measured by ELISA. No stimuli: medium control. *S.s* PRM: peptidorhamnomannan of *S. schenckii. S.b* PRM: peptidorhamnomannan of *S. brasiliensis*. The data were expressed as mean ± SEM. Statistical analysis was performed by Mann-Whitney-U test. *p <0.05, **p <0.01, ***p <0,001, ****p < 0.0001; differs from no stimuli control, or ^#^p < 0.05, ^##^p < 0.01, ^###^p < 0,001, ^####^p < 0.0001; differs between stimulated conditions.

In order to study whether PRM of both *S. brasiliensis* and *S. schenckii* are able to induce innate immune-derived cytokines, human PBMCs were stimulated with different concentrations of *S.b* PRM and *S.s* PRM. We observed that the concentrations of 10 and 100 μg/ml of *S.b* PRM were able to induce higher IL-1β, TNF-α and IL-6 levels than *S.s* PRM (**Figure 1 E, F** and **G**). Of importance, similar to *S. schenckii* yeasts, *S.s* PRM induced higher levels of the IL-1Ra when compared to *S.b* PRM (**Figure 1 I**). Both *S.b* PRM and *S.s* PRM were able to induce potent IL-8 production in comparison to unstimulated cells, however *S.b* PRM induced higher amounts of IL-8 than *S.s* PRM (**Figure 1 H**). Importantly, the PRMs of both fungi were free of residual lipopolysaccharide (LPS), since the pre-incubation of PBMCs with or without polymyxin B did not interfere in the production of IL-1β and IL-8 (**Figure S1 B** and **C**). As a positive control, IL-1β production induced by LPS was decreased in the presence of polymyxin B (**Figure S1 A**).

### *S. brasiliensis* and *S.b* PRM induced higher production of T helper cytokines than *S. schenckii* and *S.s* PRM

In order to investigate whether PRMs can induce T-lymphocyte-derived cytokines, we next evaluated the production of IFN-γ, IL-17 and IL-22 in human PBMCs after 7 days exposure with *S.s* and *S.b* PRM, as well as *S. schenckii* and *C. albicans* yeasts as control stimuli. The exposure of human PBMCs to *S. brasiliensis* induced the production of IFN-γ, IL-17 and IL-22 at comparable levels to *C. albicans* (**Figure 2 A, B** and **C**). *S. schenckii* also induced significantly the production of these cytokines, however at lower concentrations when compared to *S. brasiliensis* and *C. albicans* (**Figure 2 A, B** and **C**). Similarly, *S.b* PRM was more potent to induce IFN-γ, IL-17 and IL-22 compared to *S.s* PRM (**Figure 2 D, E** and **F**).

**Figure 2.**
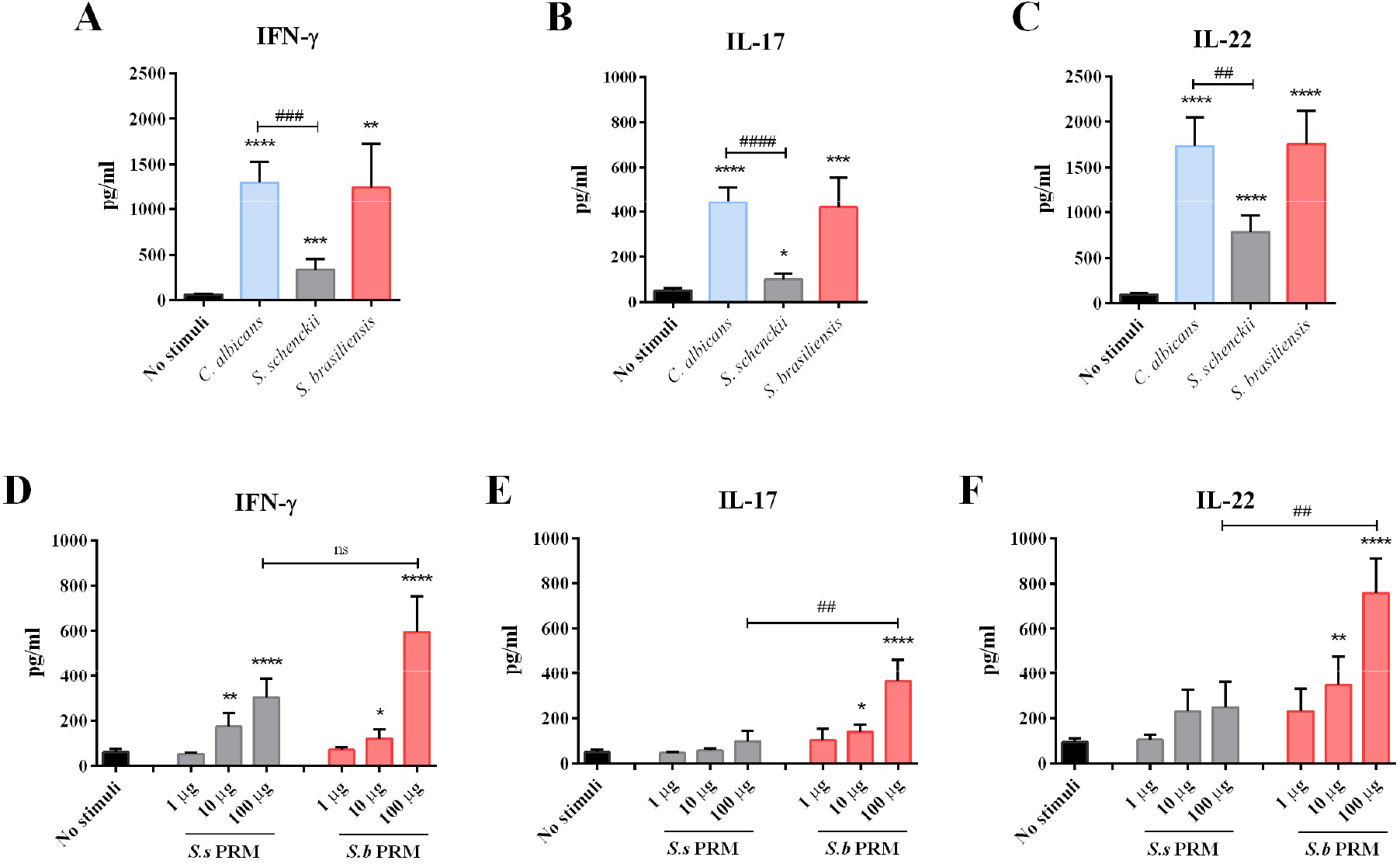
*S. brasiliensis* and *S.b* PRM strongly induce Th1 and Th17 response while *S. schenckii* and *S.s* PRM induce Th17 response marked by IL-22 production. PBMCs were stimulated with heat-killed *Candida albicans, S. schenckii* and *S. brasiliensis*. After 7 days of stimulation, supernatants were collected and **A.** IFN-γ (n=8), **B.** IL-17 (n=8) and **C.** IL-22 (n=8) production were measured by ELISA. Peptidorhamnomannans (PRMs) were evaluated at different concentrations of 1, 10 and 100 μg/ml. After 7 days of stimulation, **D.** IFN-γ (n=8), **E.** IL-17 (n=8) and **F.** IL-22 (n=8) production were measured by ELISA. No stimuli: PBMCs under the same conditions but without fungi or PRMs. The data were expressed as mean ± SEM. Statistical analysis was performed by Mann-Whitney-U test. *p < 0.05, **p < 0.01, ***p <0,001, ****p < 0.0001; differs from no stimuli control, or ^##^p < 0.01, ^###^p < 0,001, ^####^p < 0.0001; differs between stimulated conditions. ns: no significant.

### Dectin-1, CR3 and distinct TLR receptors are required for the peptidorhamnomannan-induced cytokine response

The recognition of the fungus through pattern recognition receptors (PRRs) is one of the first steps to initiate the immune response against the pathogen, leading to cytokine induction (Hardison and Brown, 2012). Dectin-1 receptor block significantly reduced the production of IL-1β and TNF-α after *S. schenckii, S.b* PRM and *S.s* PRM stimulation (**Figure 3 A, B** and **Figure S2 A, B**). Mincle receptor blockade decreased IL-1β and TNF-α production only for *S. schenckii* and *S. brasiliensis* yeasts (**Figure 3 A** and **Figure S2 A**). Blocking Mincle or Dectin-1 resulted in an increase in IL-1Ra production by cells stimulated with *S. schenckii* or *S. brasiliensis* yeasts (**Figure S2 C**). Inhibition of spleen tyrosine kinase (Syk) and noncanonical serine-threonine kinase Raf-1 resulted in a significant decrease in the production of IL-1β and TNF-α by both fungi and their respective PRMs (**Figure 3 B** and **Figure S2 B**). There was no effect in cytokine production after using anti-dectin-2 blocking antibodies. *C. albicans* was used as a control in all blocking experiments and IL-1β production by the fungus was influenced by Mincle and Dectin-1 blocking, via Syk and Raf-1 (**Figure S3 A**).

**Figure 3.**
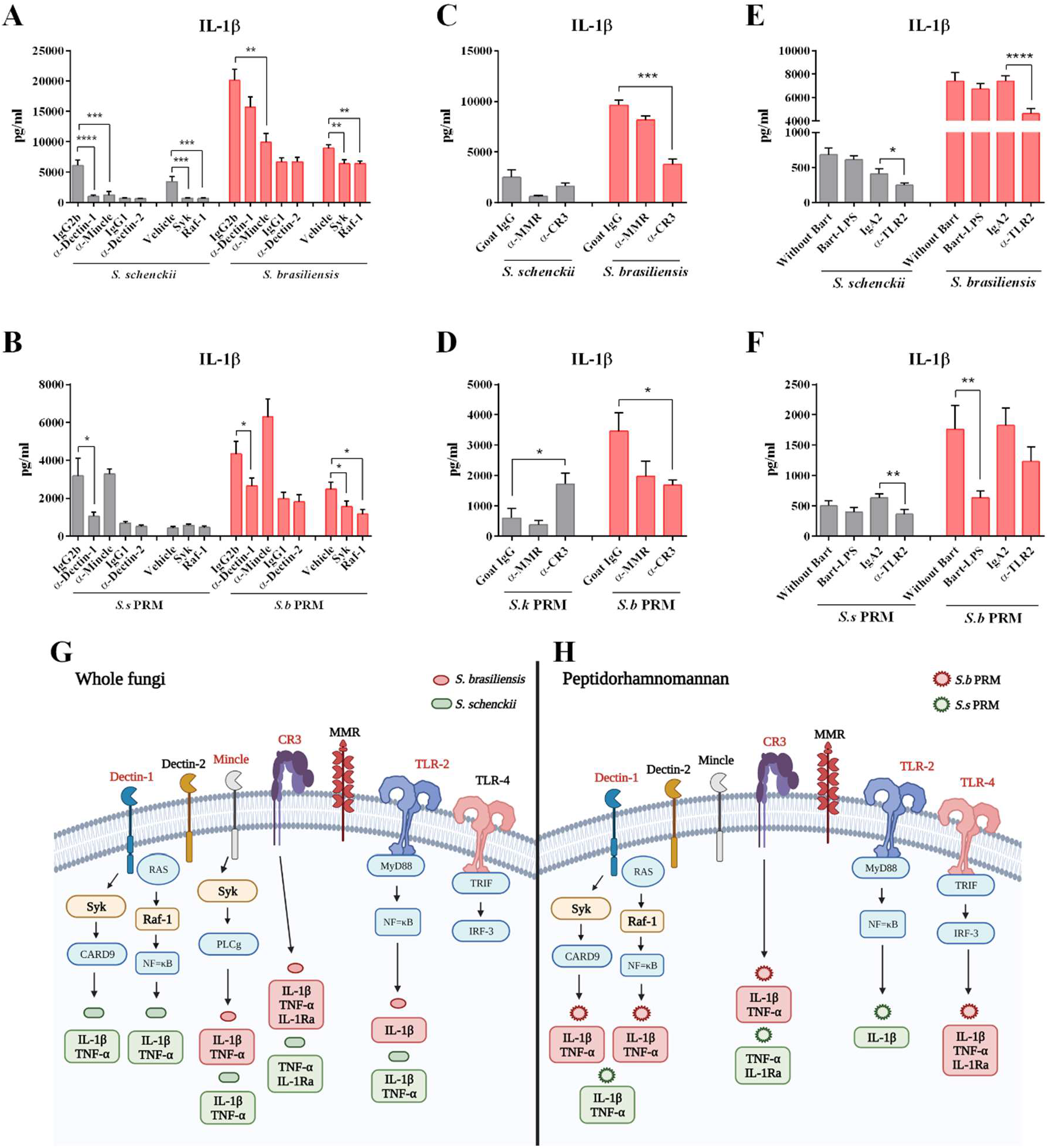
C-type lectin, TLRs receptors and signalling pathways involved in the IL-1β production by *S. schenckii, S. brasiliensis* and their respective peptidorhamnomannans. PBMCs were pre-incubated for 1 h with **A, B.** 10 μg/ml anti-dectin-1 (n=6), anti-dectin-2 (n=7) and anti-mincle (n=4) antibodies, or control isotypes IgG2b (n=6) and IgG1 (n=7). For assessing the signalling pathways, the cells were pre-incubated with 50 nM Syk (n=8), 1μM Raf-1 (n=8) and Vehicle (DMSO; n=8) as a control. **C, D.** anti-CR3 (n=8) and anti-MMR (n=4) antibodies, or isotype Goat IgG (n=8) as a control, **E, F.** anti-TLR2 (n=8) and 20 ng/ml *Bartonella quintana* LPS (n=8), or isotype IgA2 (n=8) and without Bart (RPMI only; n=8) as a control. After the blocking period, cells were stimulated with heat-killed *S. schenckii* and *S. brasiliensis* or *S.s* PRM and *S.b* PRM. After 24 hours of stimulation, supernatants were collected. IL-1β production was measured by ELISA. The data were expressed as mean ± SEM. Statistical analysis was performed by Wilcoxon test. *p <0.05, **p <0.01, ***p <0,001, ****p <0.0001; differs from controls of isotype antibody or Vehicle. Schematic overview of the receptors and signalling pathways required in the recognition of **G.** *S. schenckii* and *S. brasiliensis*, **H.** *S.s* PRM and *S.b* PRM.

Mannose receptor (MMR) and complement receptor 3 (CR3) are described as involved in the recognition of fungal N-linked mannan and β-glucans, respectively (Tang et al., 2018). Therefore, we investigated the involvement of these receptors in the cytokine response induced by the recognition of the whole fungus *Sporothrix spp*. or by the respective PRM. Blocking MMR did not interfere in IL-1β, TNF-α and IL-1Ra production by any of the stimuli tested (**Figure 3 and Figure S2**). In contrast, CR3 was required for the production of IL-1β, TNF-α and IL-1Ra by cells stimulated with *S. brasiliensis, S.s* PRM and *S.b* PRM (**Figure 3 and Figure S2**). Interestingly, blocking CR3 in cells stimulated with *S.s* PRM increased the concentration of IL-1β in the culture supernatant and decreased the concentrations of IL-Ra (**Figure 3 D** and **Figure S2 H**). Neither MMR nor CR3 were required in the cytokine response against *C. albicans* (**Figure S3 B**).

The importance of TLRs in cutaneous sporotrichosis has been widely explored (Rossato et al., 2019a, 2019b). In our study, blocking of TLR2 with specific antibody resulted in decreased production of IL-1β and TNF-α by *S. schenckii* and IL-1β by *S. brasiliensis* (**Figure 3 E** and **Figure S2 I**). The blockade of TLR4 with its natural antagonist *Bartonella quintana* LPS resulted in a significant decrease in IL-1β, TNF-α and IL-1Ra by cells stimulated with *S.b* PRM (**Figure 3 D** and **Figure S2 J, L**). IL-1β production in cells stimulated with *S.s* PRM was impaired by TLR2 blockade (**Figure 3 F**). Cells incubated in the presence of TLR4 antagonist and LPS as a control showed a significant decrease in IL-1β production compared to cells treated with LPS only (**Figure S3 C**). **Figures 3 G** and **H** summarize the receptors required in the cytokine response induced by *Sporothrix* yeasts and their PRMs.

### IL-23 drives Th17 response to *S. brasiliensis*, but not *S.b* PRM

As Th17 is very important for the host defense against fungi, we investigate the role of IL-23 on the Th17 profile. *S. brasiliensis* and its respective PRM, induced the production of IL-23 (**Figure S4 A**). To investigate the involvement of IL-23 in driving the Th17 response, we blocked the IL-23 receptor with a specific anti-IL-23 antibody. IL-17 production by *C. albicans* control was significantly decreased (**Figure S4 B** and **C**). Interestingly, blocking IL-23 decreased the production of IL-17 and IL-22 in cells stimulated with *S. brasiliensis*, but not with *S.b* PRM (**Figure S4 D, E, F** and **G**).

### The inflammatory response induced by *S. brasiliensis* and its peptidorhamnomannan is strongly dependent of IL-1

We investigated the role of IL-1 in the inflammation induced by *Sporothrix* yeasts and PRMs. Anakinra, a recombinant form of the human IL-1 receptor antagonist (IL-1 Ra), competes with IL-1β for binding to the IL-1 receptor (Cavalli and Dinarello, 2018) (**Figure 4 A**). In our study, we blocked the IL-1 receptor of human PBMCs using anakinra, followed by stimulation with *Sporothrix* yeasts and their respective PRMs. IL-1 blockade resulted in a significant decrease of IL-1β, TNF-α and IL-6 by cells stimulated with *S. schenckii, S. brasiliensis, S.s* PRM and *S.b* PRM (**Figure 4 B** and **D**). Furthermore, IL-1 blockade strongly decreased the concentration of T helper cytokines IL-17, IL-22 and IFN-γ by cells stimulated with *S. brasiliensis, S. schenckii* and *S.b* PRM (**Figure 4 C** and **E**). Regarding the *S.s* PRM, IL-1 blockade led to a decrease in the concentration of IL-17 and IL-22, while IFN-γ remains similar to the control without anakinra (**Figure 4 E**). The production of IL-1β, IL-17 and IFN-γ was significantly affected by IL-1 blockade receptor in cells stimulated with *C. albicans* control (**Figure S5 A, B** and **C**).

**Figure 4.**
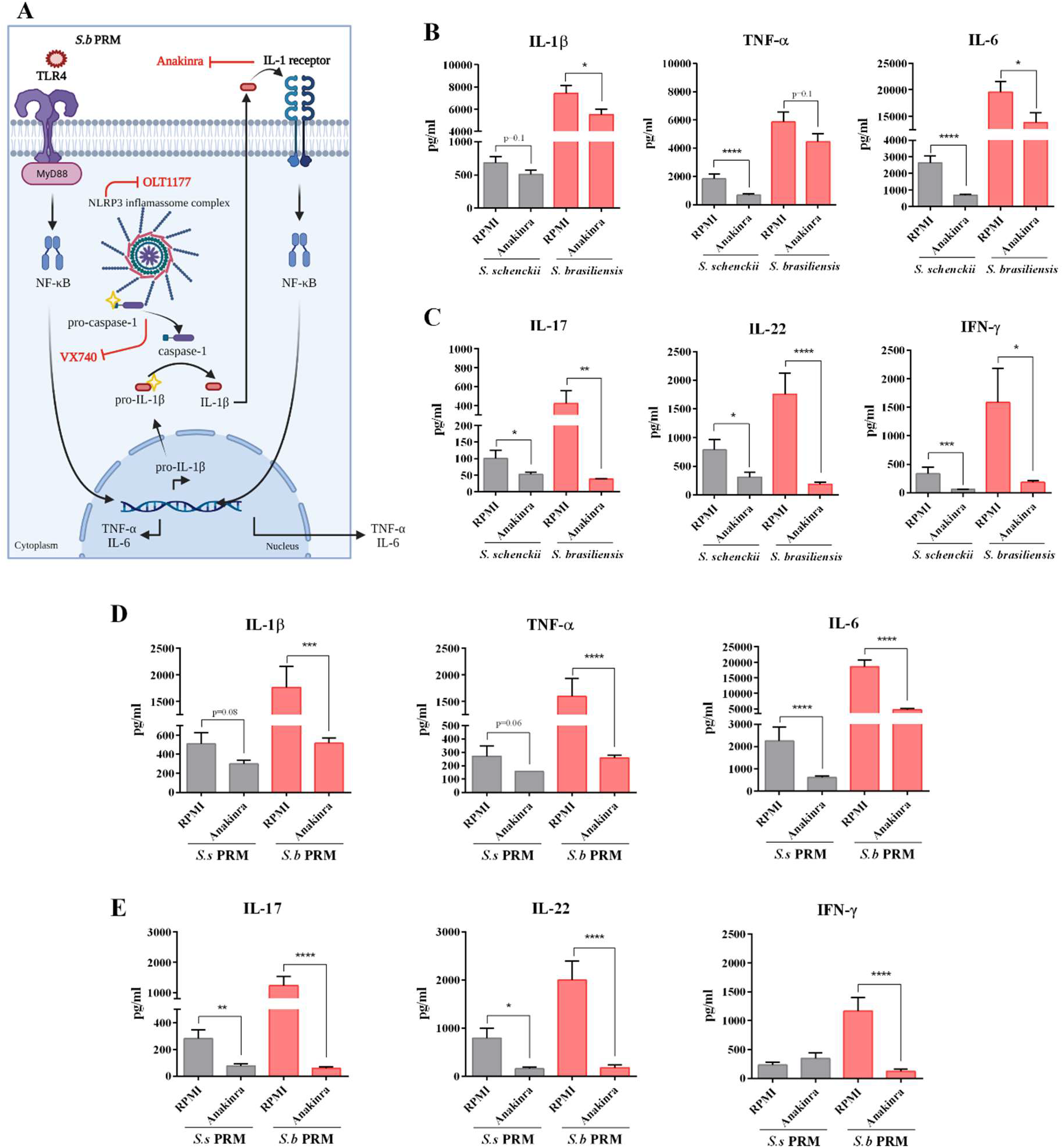
IL-1 receptor blockade significantly reduces the production of pro-inflammatory cytokines by *S. schenckii, S. brasiliensis* and their respective PRMs. **A.** Schematic overview of the inhibitors used to investigate the influence of IL-1 signalling on the inflammatory response induced by *S. schenckii, S. brasiliensis* and their respective PRMs. PBMCs were pre-incubated for 1 h with and without 10 μg/ml Anakinra. After, cells were stimulated with heat-killed *S. schenckii* and *S. brasiliensis* or *S.s* PRM and *S.b* PRM. Cells were incubated for 24 hours and 7 days at 37 °C and 5% CO_2_. After 24 hours of stimulation, supernatants were collected and IL-1β, TNF-α and IL-Ra (n=8) production were measured by ELISA in cells stimulated with **B.** *S. brasiliensis* and *S. schenckii* and **D.** *S.s* PRM and *S.b* PRM. After 7 days of stimulation, supernatants were collected and IL-17, IL-22 and IFN-γ (n=9) production were measured by ELISA in cells stimulated with **C.** *S. brasiliensis* and *S. schenckii* and **E.** *S.s* PRM and *S.b* PRM. The data were expressed as mean ± SEM. Statistical analysis was performed by Wilcoxon test. *p <0.05, **p <0.01, ***p < 0.001 ****p <0,0001; differs from RPMI control (PBMCs without anakinra and treated with the respective stimulus).

### *Sporothrix brasiliensis* and *S.b* PRM-induced inflammation dependent of caspase-1 or inflammasome activation

The production of the pro-inflammatory cytokines IL-α and IL-1β is induced by the recognition of the pathogen via the Toll-like receptor and by the subsequent activation of NF-κB. At the same time, activation of the inflammasome by other stimuli results in proteolytic cleavage of pro-IL-1β into its mature form by the action of the cysteine protease caspase-1 (Fettelschoss et al., 2011) (**Figure 5 A**). OLT1177, a β-sulfonyl nitrile compound, inhibits the NLRP3 inflammasome activation and Pralnacasan (VX740), a selective non-peptide, inhibits the conversion of pro-caspase-1 to its functional form, preventing the production and secretion of mature IL-1β (Marchetti et al., 2018; Rudolphi et al., 2003). As we found that inflammation induced by the whole fungi and their PRMs is strongly dependent on IL-1, we investigated the potential of these two compounds in controlling inflammation induced by *Sporothrix* yeasts and their PRMs.

**Figure 5.**
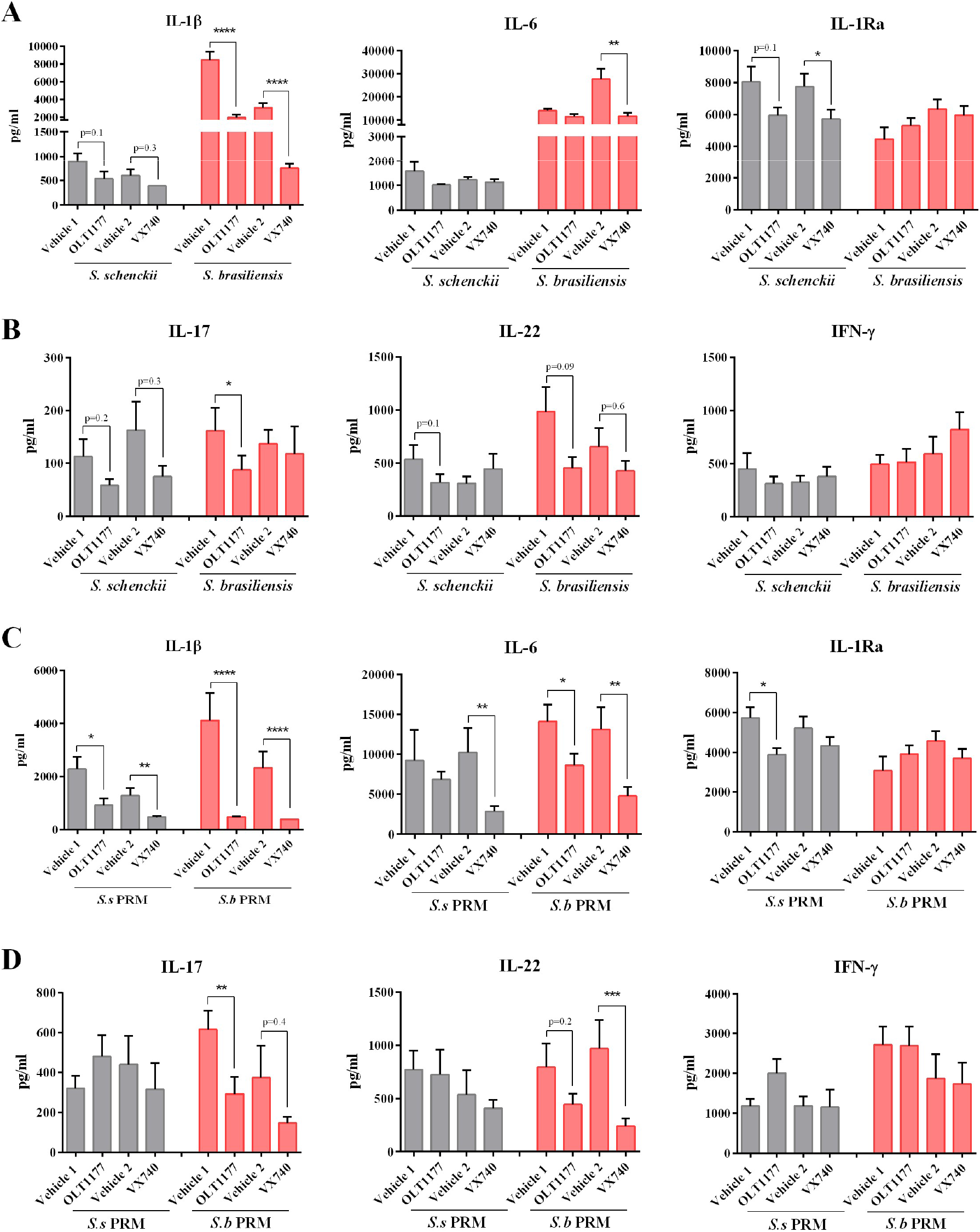
Blocking inflammasome activation by OLT1177 and caspase-1 by Pralnacasan (VX-740) results in decreased of Th17 cytokines. PBMCs were pre-incubated for overnight with 10 μM (10^3^ nM) of OLT1177 or Pralnacasan (VX-740) and their respective vehicles as controls (Vehicle 1: PBS; Vehicle 2: DMSO). After the blocking period, cells were stimulated with heat-killed *S. schenckii* and *S. brasiliensis* or *S.s* PRM and *S.b* PRM. Cells were incubated for 24 hours and 7 days at 37 °C and 5% CO_2_. After 24 hours of stimulation, supernatants were collected and IL-1β, IL-6 and IL-Ra (n=6) production were measured by ELISA in cells stimulated with **A.** *S. brasiliensis* and *S. schenckii* and **C.** *S.k* PRM and *S.b* PRM. After 7 days of stimulation, supernatants were collected and IL-17, IL-22 and IFN-γ (n=6) production were measured by ELISA in cells stimulated with **B.** *S. brasiliensis* and *S. schenckii* and **D.** *S.k* PRM and *S.b* PRM. The data were expressed as mean ± SEM. Statistical analysis was performed by Wilcoxon test. *p <0.05, **p <0.01, ***p <0.001 ****p <0,0001; differs from the treatment with vehicle control.

First, OLT1177 was evaluated at concentrations of 0.1 - 10^4^ nM followed by LPS stimulation. The compound did not show toxicity at any concentrations, and it was effective in decreasing the production of IL-1β between 100 to 10^4^ nM (**Figure S6 A** and **B**). Furthermore, the blockade did not interfere with the production of pro-IL-1β and TNF-α after stimulation with LPS (**Figure S6 C** and **D**). The blockade of NLRP3 inflammasome with 10^3^ nM of OLT1177 caused a decrease in IL-1β, in cells stimulated with *S. brasiliensis* (**Figure 5 A**), and IL-1β and IL-6 in cells stimulated with *S.b* PRM (**Figure 5 C**). IL-17 and IL-22 production was decreased in cells stimulated with both *S. brasiliensis* and *S.b* PRM (**Figure 5 B** and **D**). Inhibition of caspase-1 significantly affected the production of IL-1β and IL-6 in cells stimulated with *S. brasiliensis*, as well as the decrease of T helper-derived IL-22 (**Figure 5 A** and **B**). In case of stimulation with *S.b* PRM the effects on IL-β and IL-6 were similar to the *Sporothrix* yeasts (**Figure 5 C** and **D**).

In relation to *S. schenckii*, inflammasome inhibition decreased the production of IL-1β, IL-1Ra, as well as IL-17 and IL-22 to a lesser extent (**Figure 5 A** and **B**). The effect of *S.s* PRM was impaired only regarding to the production of cytokines associated with innate immunity such as IL-1β and IL-1Ra by blocking the inflammasome and IL-1β and IL-6 by blocking caspase-1 (**Figure 5 C** and **D**).

Neither OLT1177 nor VX740 were able to interfere with the IFN-γ induced by the whole fungus or PRMs (**Figure 5 B** and **D**). Thus, we measured IL-18 concentrations in 24-hour supernatants and observed that its production remains unchanged between conditions treated or not with the OLT1177 or VX740 inhibitors (**Figure 6 A**). LPS was used as a control in all experiments and IL-β levels were significantly decreased after blocking with OLT1177 and VX740 (**Figure 6 B**). The treatment of cells with OLT1177 and VX740 did not interfere with the production of intracellular pro-IL-1β and expression of IL-1β and TNF-α mRNA after stimulation with PRMs (**Figure 6 B, C and D**). Both compounds did not demonstrate significant toxicity in PBMCs by the proposed incubation periods in this study of 24 and 7 days as there were no differences in LDH production (**Figure 6 E**). LPS was used as a control in all experiments and IL-β levels were significantly decreased after blocking with OLT1177 and VX740 (**Figure S6 E**).

**Figure 6.**
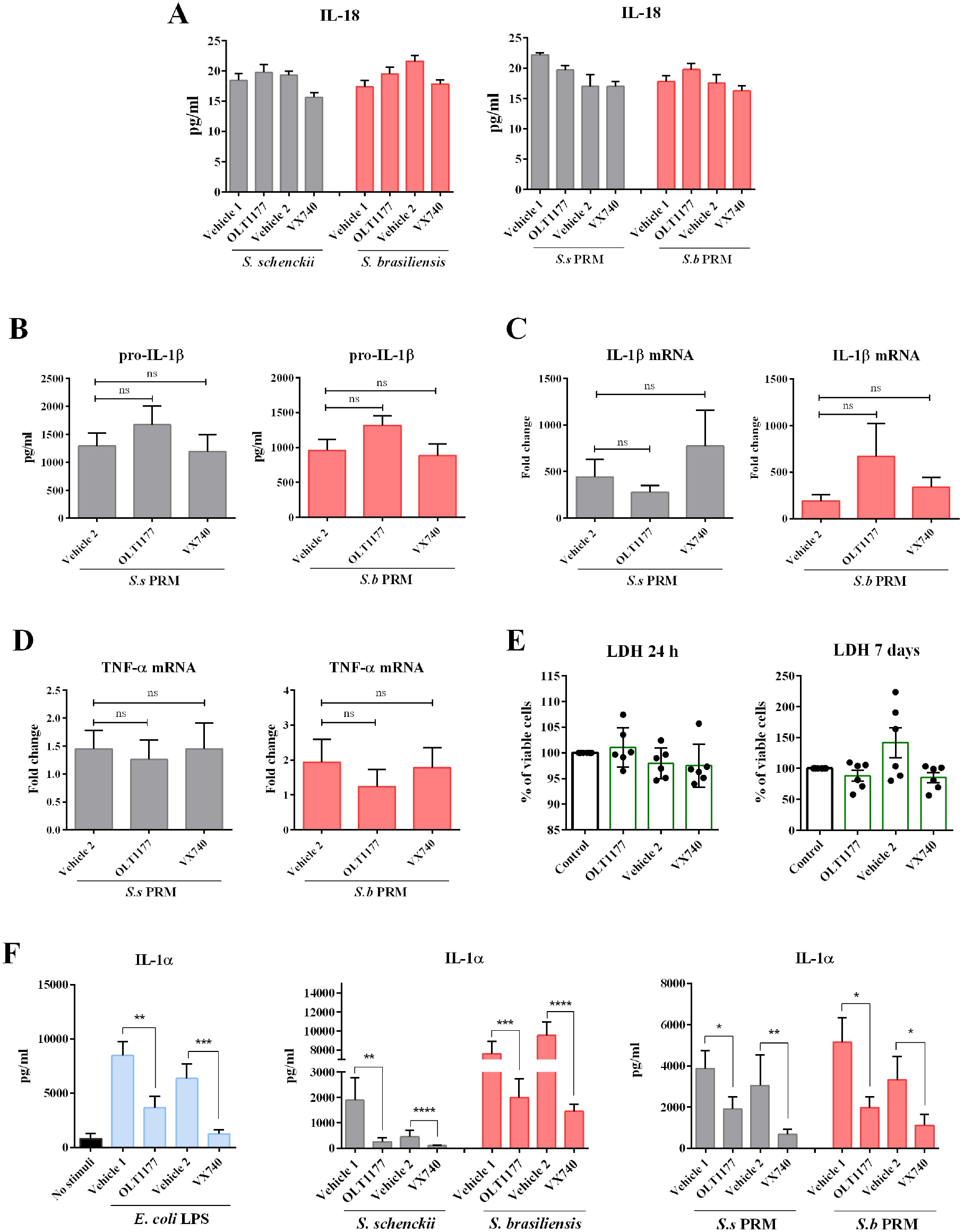
OLT1177 and VX740 exclusively affect IL-1β and IL-1α production. PBMCs were pre-incubated for overnight with and without 10 μM of OLT1177 or Pralnacasan (VX740) and their respective vehicles as controls (Vehicle 1: PBS; Vehicle 2: DMSO) (n=6). After the blocking period, cells were stimulated with LPS as a stimuli control, heat-killed *S. schenckii* and *S. brasiliensis* or *S.s* PRM and *S.b* PRM. After 24 hours of stimulation, supernatants were collected to measure **A.** IL-18 and **B.** The cells were lysed and the intracellular IL-1β production were measured by ELISA. Expression of **C.** IL-1β and **D.** TNF-α mRNA by real-time qPCR. **E.** Percentage of cell viability after 24 hours and 7 days in the presence of OLT1177 and VX740. **F.** Cells were lysed and the intracellular IL-1α (n=6) concentration was measured by ELISA. Statistical analysis was performed by Wilcoxon test. *p < 0.05, **p < 0.01, ***p < 0.001; differs from the treatment with vehicle control.

The effect of OLT1177 and VX740 inhibitors on the production of cytokines IL-6, IFN-γ, IL-17 and IL-22 by *Sporothrix* yeasts or PRMs was less significant than Anakinra. Therefore, we decided to investigate whether this moderate effect could be due to IL-1α expression, since blockade of the IL-1 receptor by anakinra prevents the binding of both IL-β and IL-1α to the receptor. Interestingly, IL-1α concentrations were decreased by inhibition of the inflammasome after stimulation with the whole fungus and PRMs treatments (**Figure 6 F**). These results demonstrate that IL-1α expression induced by *Sporothrix* yeasts and theirs PRMs is dependent on inflammasome and caspase-1 activity.

### *Sporothrix schenckii* and *S.s* PRM induces IL-1β independently of reactive oxygen species

Inflammasome activation may be mediated by reactive oxygen species (ROS) (Spooner and Yilmaz, 2011). To investigate whether IL-1β production was associated with inflammasome activation by ROS we measured the release of ROS produced during 1 hour after the addition of *Sporothrix* yeasts and PRMs. Zymosan was used as a positive control for ROS production. None of the whole fungi or PRM were able to induce ROS production similarly to zymosan (**Figure 7 A**). *S. brasiliensis* induced the higher production of IL-1β than *S. schenckii* (**Figure 7 C**). *S.s* PRM and *S. schenckii* induced the production of ROS and IL-1β at low concentrations when compared to zymosan, *S. brasiliensis* and *S.b* PRM (**Figure 7 B** and **C**). Interestingly, *S.b* PRM did not induce significant amounts of ROS compared to the unstimulated control, but it induced higher concentrations of IL-β than *S.s* PRM (**Figure 7 B** and **C**). To investigate whether IL-1β induced by *S.b* PRM was completely independent of ROS, we blocked ROS production using diphenyleneiodonium chloride (DPI), an inhibitor of NADPH oxidase. DPI was able to significantly reduce ROS production in cells stimulated with *Sporothrix* yeasts, PRMs and zymosan as a control (**Figure 7 D**). After 24h, IL-1β production in cells stimulated by *S. brasiliensis* and *S.b* PRM was significantly decreased by low ROS production (**Figure 7 E**). Surprisingly, *S. schenkii* and S.s PRM did not have modify their IL-1β production if ROS release was inhibited (**Figure 7 E**). ROS production after 24h remained around 4×10^3^ RLU, same values observed for non-stimulated controls (**Figure S7**).

**Figure 7.**
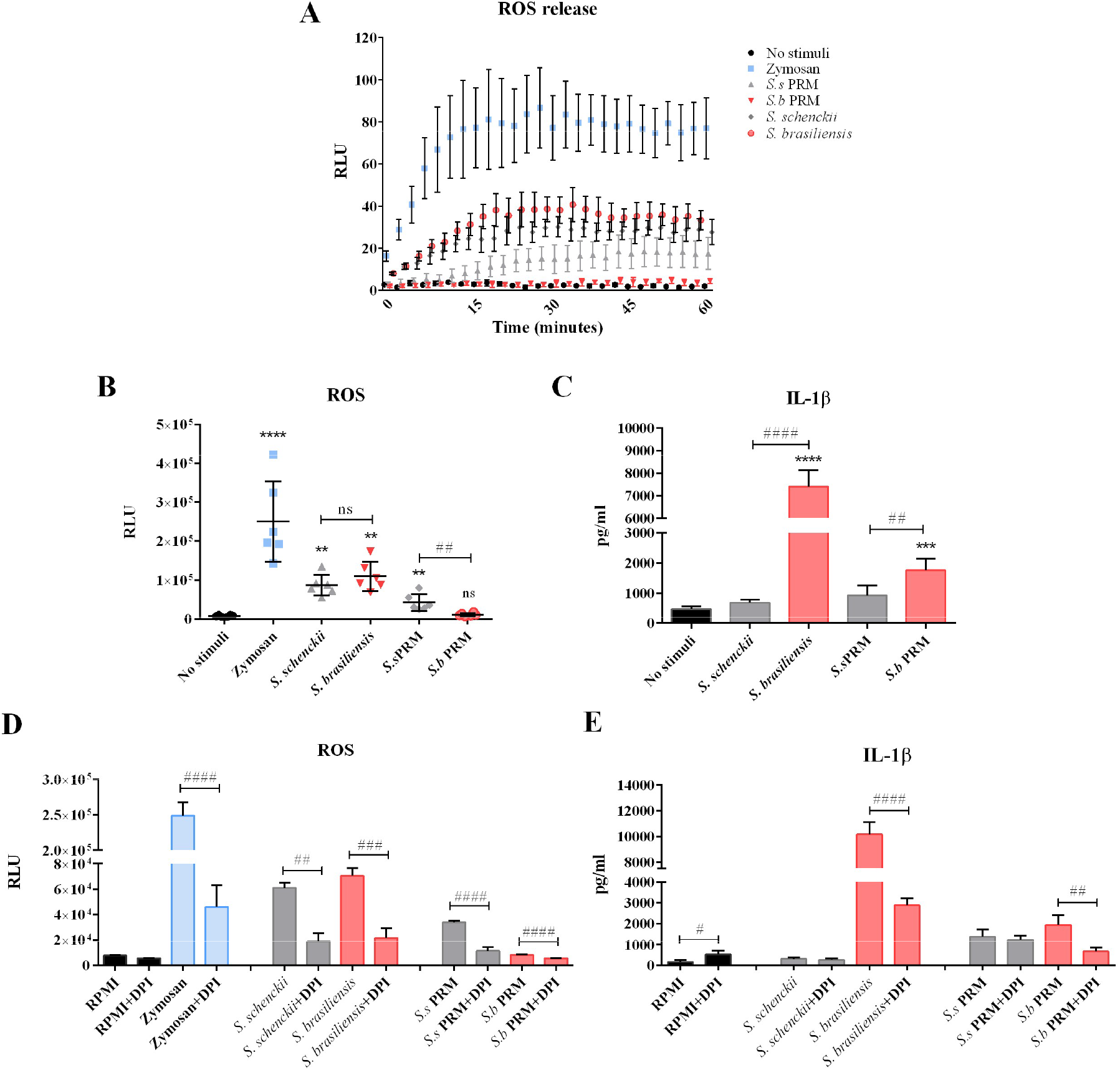
Relative ROS production induced by *S. schenckii, S. brasiliensis* and their respective PRMs. **A.** Kinetics of ROS release by PBMCs stimulated with zymosan, *S.s* PRM, *S.b* PRM and heat-killed *S. schenckii* and *S. brasiliensis* over a period of 1 hour. **B.** Total ROS production measured by area under each curve for the whole period of 1 hour. ROS production was measured by luminol-enhanced chemiluminescence assay **C.** IL-1β production after 24 hours of stimuli was measure by ELISA. Cells were treated with 10 μM diphenyleneiodonium chloride (DPI) for 30 min and stimulated with Zymosan, *Sporothrix* yeast and PRMs. **D.** Total ROS production measured by area under each curve for the whole period of 1 hour. **E.** IL-1β production after 24 hours of stimuli was measure by ELISA. The data were expressed as mean ± SEM. Statistical analysis was performed by Wilcoxon test. **p < 0.01, ***p <0,001, ****p < 0.0001; differs from no stimuli control, or ^##^p < 0.01, ^####^p < 0.0001; differs between stimulated conditions.

## Discussion

In the present study, we identify the mechanisms involved in the pattern recognition and induction of cytokines by PRMs, cell wall glycoconjugates of *Sporothrix spp*. Recently identified as an cell wall pathogen associated molecular pattern (Neves et al., 2021). Furthermore, we identified the pattern recognition receptors of human PBMCs responsible for the induction of IL-1β, TNF-α and IL-1Ra by PRMs and yeasts from *S. brasiliensis* and *S. schenckii*. We demonstrate that *S. brasiliensis* and *S.b* PRM are capable to induce a robust inflammatory response that is strongly IL-1-dependent, while *S. schenckii* and *S.s* PRM induce the production of higher amounts of IL-1Ra. These results suggest that IL-1β is a crucial player in the regulation of the immune response induced by *S. brasiliensis* and improve our understanding about the pathogenicity of *S. schenckii*. In addition, these findings open novel treatment strategies towards the control of exacerbate inflammation induced by *S. brasiliensis* infection in humans.

Glycoconjugates present in the cell wall of pathogenic fungi are known to stimulate or suppress the host’s immune response (Erwig and Gow, 2016). PRMs are a major glycopeptide component of the cell wall of *S. schenckii* and *S. brasiliensis*, and it is composed by N- and O-glycan chains linked to a peptide moiety (Lopes-Bezerra, 2011). The PRMs are located in the external microfibrillar layer of the fungus cell wall, while structural polysaccharides, chitin and β-glucan molecules, are present in the inner cell wall layer (Lopes-Bezerra et al., 2018b). Because of its location, the PRM was shown to be the fungus cell wall component that firstly interact with the host immune cells and consequently initiate the immune response (Lopes-Bezerra et al., 2018b; Neves et al., 2021). Our results show that the cytokine profiles induced in PBMCs *in vitro* was similar for *S. brasiliensis* yeasts and *S.b* PRM, with high production of pro-inflammatory cytokines. On the other hand, *S. schenckii* and *S.s* PRM induced high levels of IL-1Ra.

Of particular interest, the immune response induced among *S. brasiliensis* and *S. schenckii* or *S.b* and *S.s* PRM are distinct. In general, sporotrichosis is manifested in several clinical forms, and the most frequent are the fixed cutaneous, lymphocutaneous and disseminated cutaneous (Queiroz-Telles et al., 2019). A retrospective study in Venezuela analysed patient isolates and discovery that infections caused by *S. schenckii* were related to the lymphocutaneous form (Camacho et al., 2015). In contrast, *S. brasiliensis* has high expression of virulence factors, in addition to being more commonly associated with disseminated manifestations of the disease (Almeida-Paes et al., 2015; Alves et al., 2020; Orofino-Costa et al., 2013; Silva-Vergara et al., 2012). In our study, stimulation of PBMCs with *S. brasiliensis* led to a more pronounced inflammatory response than *S. schenckii*. This result corroborates the findings in the literature in which we can relate the higher level of pathogenicity reported in infections by *S. brasiliensis* with the greater capacity to generate an exacerbated inflammatory response in the host, that increases tissue destruction and results in aggravation of the disease (Morgado et al., 2018).

We also report potent release of IL-1Ra by the immune cells stimulated with *S. schenckii*. IL-1β and IL-1α are pro-inflammatory cytokines that are normally elevated after traumatic injuries and infections (Vuolteenaho et al., 2003). These cytokines are responsible for the induction of fever, inflammation, tissue destruction and in severe cases, septic shock. In this way, IL-1 axis is a potent pro-inflammatory pathway that is regulated by IL-1Ra. IL-1Ra can inhibit pathway activation by binding to the IL-1 receptor type I on the surface of the host cell and thus control inflammation induced by both IL-1β and IL-1α (Cavalli and Dinarello, 2018; Vuolteenaho et al., 2003). Suppression of the anti-inflammatory cytokine IL-1Ra signalling pathway was studied in invasive aspergillosis (Gresnigt et al., 2014). The authors demonstrated that the complete absence of IL-1Ra protects mice from developing invasive aspergillosis. In contrast, the production of high levels of IL-1Ra results in increased susceptibility to infection (Gresnigt et al., 2014). We observed that *S. schenckii* and its PRM induce high levels of IL-1Ra. Therefore, our findings suggest that severe cases of sporotrichosis caused by *S. schenckii* may be associated with high levels of IL-1Ra induced by the fungus’s PRM, that inhibit the IL-1 pathway as an important component of the host defense.

In the present study, dectin-1 receptor and its downstream signalling molecules Syk and Raf-1 were required for the production of IL-1β and TNF-α induced by *S. schenckii* yeasts only. The importance of dectin-1 for the recognition of *S. schenckii* by PBMCs has been previously reported, however, the authors attributed to the recognition of β-1,3-glucan (Martínez-Álvarez et al., 2017). Here we demonstrate that dectin-1 also plays a role in the cytokine induction by *S.s* PRM. Furthermore, we demonstrate that the Mincle receptor is required for cytokine production induced by *S. schenckii* and *S. brasiliensis* yeasts, but is not involved in the PRM response of either species. These results suggest that the Mincle receptor may recognize other components of the *Sporothrix* spp. cell wall that are important in the human response against the yeast cells. However, which are the PAMPs that the Mincle receptors are able to recognize in fungi cell wall remains to be elucidated. To date, studies report that Mincle recognizes α-mannose, glyceroglycolipid and mannosyl fatty acids linked to mannitol in *Malassezia* spp. (Goyal et al., 2018; Yamasaki et al., 2009). Mannosylinositolphosphorylceramides, glycolipids bearing α-mannose, were already described on the cell surface of the yeast form of *S. schenckii* (Loureiro y Penha et al., 2001).

Among pattern recognition receptors, TLRs are the most explored and their importance to cutaneous sporotrichosis has already been demonstrated (Rossato et al., 2019a, 2019b). In this study, we demonstrate in human PBMCs that TLR-2 and TLR-4 participate in the cytokine response induced by PRM of *S. schenckii* and *S. brasiliensis*, respectively. In addition, CR3 proved to be a crucial receptor, as it was required for the recognition of *S. brasiliensis*, *S. schenckii* and its respective PRMs. Our data corroborates a recent study in human monocyte derived macrophages that have demonstrated that PRM of *Sporothrix* spp. is recognized by CR3 receptor component. Interestingly, when both CR3 and TLR-4 are blocked a clear cooperation between these two receptors was observed with the secretion of a soluble PRR, PTX3 (Neves et al., 2021).

The Th17 response is crucial for host protection during sporotrichosis (Verdan et al., 2012). IL-23 is a cytokine with pro-inflammatory properties, and is crucial for the induction of IL-17. The role of the IL-23/IL-17 axis in inflammation has been demonstrated for autoimmune diseases and candidiasis (Nur et al., 2019; Tang et al., 2012). In our study, we demonstrated that only *S. brasiliensis* yeasts and *S.b* PRM induce significant amounts of IL-23 in human PBMCs. Interestingly, the treatment of cells with anti-hIL-23 affected the IL-17 and IL-22 induction only upon *S. brasiliensis* yeasts exposure. This result suggests that IL-23 is redundant for the induction of a Th17 response by *S.b* PRM. A recent study demonstrated a new role for IL-23 independent of IL-17 in systemic candidiasis: the authors noted that the absence of IL-23 led to massive neutrophil and monocyte death during infection, suggesting that IL-23 is also a key regulator of myeloid cell dynamics responsible for ensuring cell survival in tissues during infection (Nur et al., 2019).

The IL-1 family of cytokines are related to innate immunity and are essential for host protection against infectious agents (Dinarello, 2018). Furthermore, the importance of IL-1 in the development of Th1 and Th17 responses in other fungi such as *C. albicans* is already well known (Verma et al., 2018; Vonk et al., 2006). However, members of the IL-1 family are also closely associated with harmful inflammation if inappropriately activated (Dinarello, 2018). In our study, we demonstrated that the immune response against *S. brasiliensis* is strongly dependent on IL-1. Blocking the inflammasome, caspase-1, and especially the IL-1 receptor resulted in a decrease in innate and T-helper inflammatory cytokines. These results suggest that the control of inflammation induced by *S. brasiliensis* can be mediated by IL-1 blockade. However, immunotherapy using cytokine blockers to treat infectious diseases needs to be conducted with caution and further studies are needed to assess the benefits vs. potential harms in the context of sporotrichosis. Currently, the greatest concern in using anti-cytokine agents is to make the patient susceptible to other opportunistic infections as commonly reported in anti-TNFα therapies (Solovic et al., 2010). However, patients treated with IL-1 blocker anakinra rarely developed opportunistic infections, with cases of viral-type upper respiratory tract infections being reported in the same proportion as those caused by biological products (Cavalli and Dinarello, 2018; Dinarello, 2018). Furthermore, due to the clinical characteristics of sporotrichosis that mainly affects the cutaneous tissue, we suggest that approaches aimed at the topical application of IL-1 inhibitors may be more suitable to control inflammation and destruction of host tissue at the site of infection. We emphasize that more studies need to be carried out in animal and human models to define the type of blocker, treatment efficacy, and duration.

Interestingly, inflammasome blockade and caspase-1 were not able to significantly influence the Th17 and Th1 response induced by *S. schenckii* and *S.s* PRM, although IL-1β induced by *S.s* PRM was significantly decreased. These results suggest that the induction of the *S. schenckii* T helper response may be dependent on a different pathway. Blockade of the IL-1 receptor significantly decreased IL-6 production in cells stimulated by *S. schenckii* and *S.s* PRM. A previous study demonstrate that Th17 polarization occurs by IL-1β and can be increased by IL-6 (Acosta-Rodriguez et al., 2007). However, more studies need to be carried out to determine the role of IL-6 in the Th17 response of *S. schenckii*.

The host’s immune system is able to generate microbicidal ROS to eliminate infectious agents (Chauhan et al., 2006). In our study, stimulation of PBMCs with *Sporothrix* yeasts and its PRMs resulted in moderate to low amounts of ROS. Comparison of this result with other studies on *Sporothrix* is particularly difficult as these studies did not use positive controls for the production of ROS, such as zymosan (Kajiwara et al., 2004; Mario et al., 2017; Ortega et al., 2015). It is known that ROS production can activate the inflammasome and induce increased IL-1β production. As IL-1β proved to be crucial for the pathogenesis of *S. brasiliensis* we investigated whether ROS was essential for inflammasome activation. ROS blockade did not influence IL-1β production by *S. schenckii* and *S.s* PRM. We can conclude that IL-1β induction in *S. schenckii* stimulated PBMCs can occur independently of ROS. The induction and maturation of IL-1β could be attributed, for example, to TLRs receptors and activation of caspase-8 (Maelfait et al., 2008).

In conclusion, our results demonstrate that the PRM from the cell wall of *S. schenckii* and *S. brasiliensis* is able to modulate the innate and adaptive immune response similar to the intact fungus. In addition, we describe for the first time the pattern recognition receptors of PBMCs involved in the cytokine immune response induced by PRMs. Ours findings reporting differences of immunological responses induced by *S. schenckii* and *S. brasiliensis* contribute to understand the pathogenesis of these two species related to human and feline cases but with different outcomes. *S. brasiliensis* yeasts and *S.b* PRM induces a potent Th17 inflammatory response in an IL-1 dependent manner. Since *S. brasiliensis* is associated with more severe cases of the disease and greater tissue destruction, treatment strategies aimed at controlling local inflammation may be valuable. In this study we demonstrate that anakinra and OLT1177 can be seen as candidates for future studies as IL-1 inhibitors to control inflammation induced by *S. brasiliensis*.

## Material and Methods

### Ethics statement and Volunteers

The study was approved by Ethics Committee of Radboud University Nijmegen, The Netherlands (NL32357.091.10). Peripheral blood mononuclear cells (PBMCs) were isolated from Buffy coats from healthy donors after written informed consent (Sanquin Blood Bank, Nijmegen, the Netherlands). Samples were anonymized to safeguard donor privacy. All the experiments were conducted according to the principle expressed in the Declaration of Helsinki.

### Fungal culture and isolation of peptidorhamnomannan (PRM)

In this study, reference strains of *Sporothrix* were registered at ATCC by Dr. Lopes-Bezerra (*S. schenckii* ATCC MYA 4820; *S. brasiliensis* ATCC MYA 4823 and used in this study. The fungus was kept in a yeast phase and expanded in 500 mL of brain Heart Infusion Broth (BHI) medium for 4 days at 37°C. Next, yeast cells were centrifuged and washed 3 times with PBS pH 7.4. Yeast cells were used to obtain heat-kill yeasts and to isolate the PRMs. Cell heat-killing was achieved by incubating at 56°C for 1 h. PRM was purified by precipitation of its borate complex with hexadecyltrimethylammonium bromide as previously described (Lloyd and Bitoon, 1971). The PRM fraction was dialyzed against ultrapure water and lyophilized.

### Peripheral blood mononuclear cells (PBMCs) isolation and treatments

PBMCs were isolated from buffy coats obtained after informed consent oh healthy volunteers (Sanquin BloodBank, Nijmegen). In brief, blood was diluted in PBS and fractions were separated by Ficoll-Paque (GE healthcare, Zeist, The Netherlands) density gradient centrifugation as previously described (Domínguez-Andrés et al., 2021). Cells were washed twice with PBS and resuspended in RPMI 1640 culture medium (Roswell Park Memorial Institute medium, Invitrogen, USA) supplemented with 10 μg/ml gentamicin, 10 mM L-glutamine, and 10 mM pyruvate (Life Technologies). The cells were counted using the Sysmex hematology instrument and the concentration was adjusted to 5×10^6^/ml (or 5×10^5^ cells in 100 ul/well) in round-bottom 96-wells plates in the presence of culture medium with 10% human pool serum. The serum was obtained from a pool of healthy donors. Cells were stimulated with 100 μg/ml of *S.s* PRM and *S.b* PRM as well as 1×10^6^ cells/ml of heat-killed *Candida albicans, S. schenckii* and *S. brasiliensis* as controls. To eliminate the possibility of lipopolysaccharide (LPS) contamination in pure PRMs, the cells were treated with the stimulus as described above after pre-incubation with 2 μg/ml of polymyxin B.

### Receptor and intracellular signalling inhibition

For inhibition experiments, 5×10^6^ cells/ml PBMCs was added per well of a 96-well roundbottom cell culture plate and pre-incubated for 1 h with different inhibitors. Pattern recognition receptors were inhibited with 20 ng/ml *Bartonella quintana* LPS, 10 μg/ml of anti-dectin-1, anti-dectin-2, anti-Mincle, anti-CR3, anti-MMR and anti-TLR2 antibodies with their respective isotype control antibodies IgG2b, IgG1, Goat IgG and IgA2 (R&D Systems). For inhibition of downstream signalling pathways molecules, the cells were pre-incubated with 50 nM of Syk inhibitor (574711, EMD-Millipore) and 1μM of Raf-1 (GW5074, Sigma). To block the IL-23 cytokine receptor, 10 μg/ml of anti-hIL-23 or control isotype Rabbit IgG was used. For all experiments, cells were treated only with RPMI or when necessary, RPMI+DMSO (vehicle) were used as controls. After the blocking period, cells were stimulated with 1×10^6^ cells/ml of heat-killed *S. schenckii* and *S. brasiliensis*, 100 μg/ml of *S.s* PRM and *S.b* PRM and 10 ng/ml of LPS from *E. coli* serotype O55:B5. The cells were incubated at 37°C, 5% CO_2_. After 24 hours, the supernatants were collected to measure cytokines production.

### Blocking IL-1 production

To investigate the role of IL-1 in the inflammatory response induced by *Sporothrix*, IL-1 receptor, NLRP3 inflammasome and caspase-1 were blocked. PBMCs (5×10^6^ cells/ml) was added per well of a 96-well round-bottom cell culture plate and pre-incubated for 1 h with and without 10 μg/ml Anakinra. To block NLRP3 inflammasome and capase-1, PBMCs were preincubated for 20 hours at 37°C, and 5% CO_2_ with and without 10 μM of OLT1177 (Olatec Therapeutics) or Pralnacasan (VX740 – Vertex, USA). After the blocking period, PBMCs were stimulated with from *S.b* PRM and *S.s* PRM (100 μg/ml) as well as heat-killed yeasts of *S. brasiliensis, S. schenckii* (1×10^5^ cells/well) and LPS (10 ng/ml) as control stimulus. Cells were incubated for 24 hours and 7 days at 37 °C and 5% CO_2_.

### Lactate dehydrogenase (LDH) measurement

Cell death was monitored using CytoTox 96 Non-radioactive cytotoxicity assay (Promega). PBMCs were treated with OLT1177, VX740 and DMSO as described above. After 24 h and 7 days of treatment, 50 μl of supernatant from all wells were transfer to a fresh 96-well flatbottom cell culture plate. Substrate mix was diluted in 12 ml of assay buffer in the absence of light and 50 μl of reconstituted substrate mix was added to each well containing the supernatants. The plate was incubated in the dark for 30 min at room temperature. After the incubation time, 50 μl of stop solution was added to each well. The absorbance was measure at 490-492 nm using the BioTek Synergy HTreader.

### Reactive oxygen species (ROS) assessment

The ability to induction of ROS were measured by oxidation luminol (5-amino-2,3, dihydro-1,4-phtalazinedione). A total of 5×10^5^ cells from nine health donors was added per well of a white-96 well assay plate in a volume of 100 μl. Next, the cells were stimulated with 100 μg/ml of *S.s* PRM and *S.b* PRM and 1×10^6^ cells/ml of heat-killed *S. schenckii* and *S. brasiliensis* opsonized. Seventy-five μg/ml of serum-treated zymosan was used as a control of ROS production. Immediately after adding the stimuli, luminol in the concentration of 1 mM was diluted in hanks balanced salt solution (HBSS) containing 0,5% of bovine serum albumin (BSA) and added 20 μl to each well with their respective stimuli. Chemiluminescence was measured in BioTek Synergy HT reader. The reading was performed at 37°C for every minute during 1 h. For blocking assays, cells were treated with 10 μM diphenyleneiodonium chloride (DPI) for 30 min followed by stimulation as described above.

### Quantification of mRNA expression by quantitative real-time PCR (qPCR)

PBMCs from six healthy donors were treated with OLT1177 and VX740 overnight and stimulated as described above. After 16h, cells were lysed with RLT buffer and the RNA was subsequently extracted using the RNeasy Mini Kit (Qiagen), following the manufacturer’s protocol. Quantification and quality assessment of extracted RNA was performed using a Nanodrop 1000 spectrophotometer (Thermo Fisher Scientific). RNA input was normalized to 100 ng for all donors. mRNA was transcribed into cDNA using iScript cDNA synthesis kit (Bio-Rad, Hercules, CA, USA). qPCR was performed using power SYBR green PCR master mix (applied Biosystems, Carlsbad, CA, USA). Primers forward (F) ACAGATGAAGTGCTCCTTCCAG and reverse (R) CATGGCCACAACAACTGACG were used for amplification of human IL-1β, F TGTTGTAGCAAACCCTCAAGC and R GAGGTACAGGCCCTCTGATG TNF-α, and F ATTGAAATCAGCCAGCACG and R AGGAACCACAGTGCCAGAT RPL37A. PCR was performed using the following conditions: 2 min 50°C, 10 min 95°C followed by 40 cycles at 95°C for 15 sec and 60°C for 1 min (applied Biosystems 7300 real-time PCR system). The RNA analysis was normalized against the housekeeping gene RPL37A calculating delta delta Ct [ΔΔCt = ΔCt (treated sample) – ΔCt (untreated sample)] and fold gene expression [2^−ΔΔCt^].

### Cytokine measurements

Cytokine levels were measured with an ELISA assay according to the manufacturer’s protocol (R&D Systems, Minneapolis, MN). IL-1β, TNF-α, IL-6, IL-8, IL-1Ra, IL-23 and IL-18 were measured after 24 h of stimulation and IFN-γ, IL-17 and IL-22 after 7 days of stimulation. IL-1α and IL-1β was measured after cell lysis with a solution of 0.5% Triton-X 100 in cells stimulated for 24 h.

### Statistical analyses

The software GraphPad Prism 6 (San Diego, CA, USA) was used for statistical analysis. The data were expressed as mean ± standard error of the mean (SEM). Differences between experimental groups were tested by Mann-Whitney *U* test or by Wilcoxon paired test as indicated. Values of \**p* ≤ 0.05, \*\**p* <0.01, \*\*\**p* < 0.001 or \*\*\*\**p* < 0.0001 were considered statistically significant. All experiments were performed on 3 independent days and at least duplicates per experiment

## Acknowledgment

Olatec Therapeutics was kindly acknowledged for supplying the inflammasome inhibitor OLT1177.

## CRediT authorship contribution statement

Brenda Kischkel: Data curation, Formal analysis, Validation, Writing - original draft. Leila Lopes-Bezerra: Conceptualization, resources, Writing - review & editing. Carlos P. Taborda: Funding acquisition, Writing - review & editing. Leo A. B. Joosten: Conceptualization, Funding acquisition, Project administration, Supervision, Writing - review & editing. Jéssica Cristina dos Santos: Conceptualization, Supervision, Writing - review & editing. Mihai G. Netea: Conceptualization, Funding acquisition, Supervision, Writing - review & editing.

