## Supplemental Figures 1-7 for "Differential recognition and cytokine induction by the peptidorhamnomannan from *Sporothrix brasiliensis* and *S. schenckii*"

### Supplementary Information - Figures

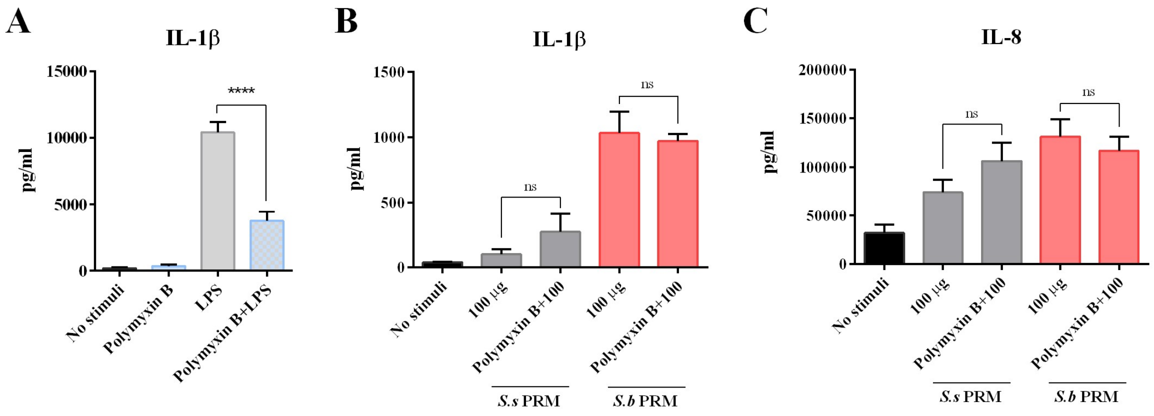


**Figure S1. PBMCs stimulation with LPS and *S.s* and *S.b* PRMs in the presence of Polymyxin B.** PBMCs were stimulated with 100 µg/ml of PRMs or 10 ng/ml of LPS for 24 hours. Polymyxin B + PRMs was used as a control to rule out the presence of LPS in pure PRMs. A. IL-1β (n=10) in PBMCs stimulated with LPS, B. IL-1β (n=5) and IL-8 (n=8) in PBMCs stimulated with PRMs. Cytokine production were measured by ELISA. ****$P < 0.0001$ Statistical analysis was performed by Mann-Whitney-U test comparing respective the treatments with RPMI as control and 100 µg/ml of PRMs with Polymyxin B + 100 µg/ml of PRMs.

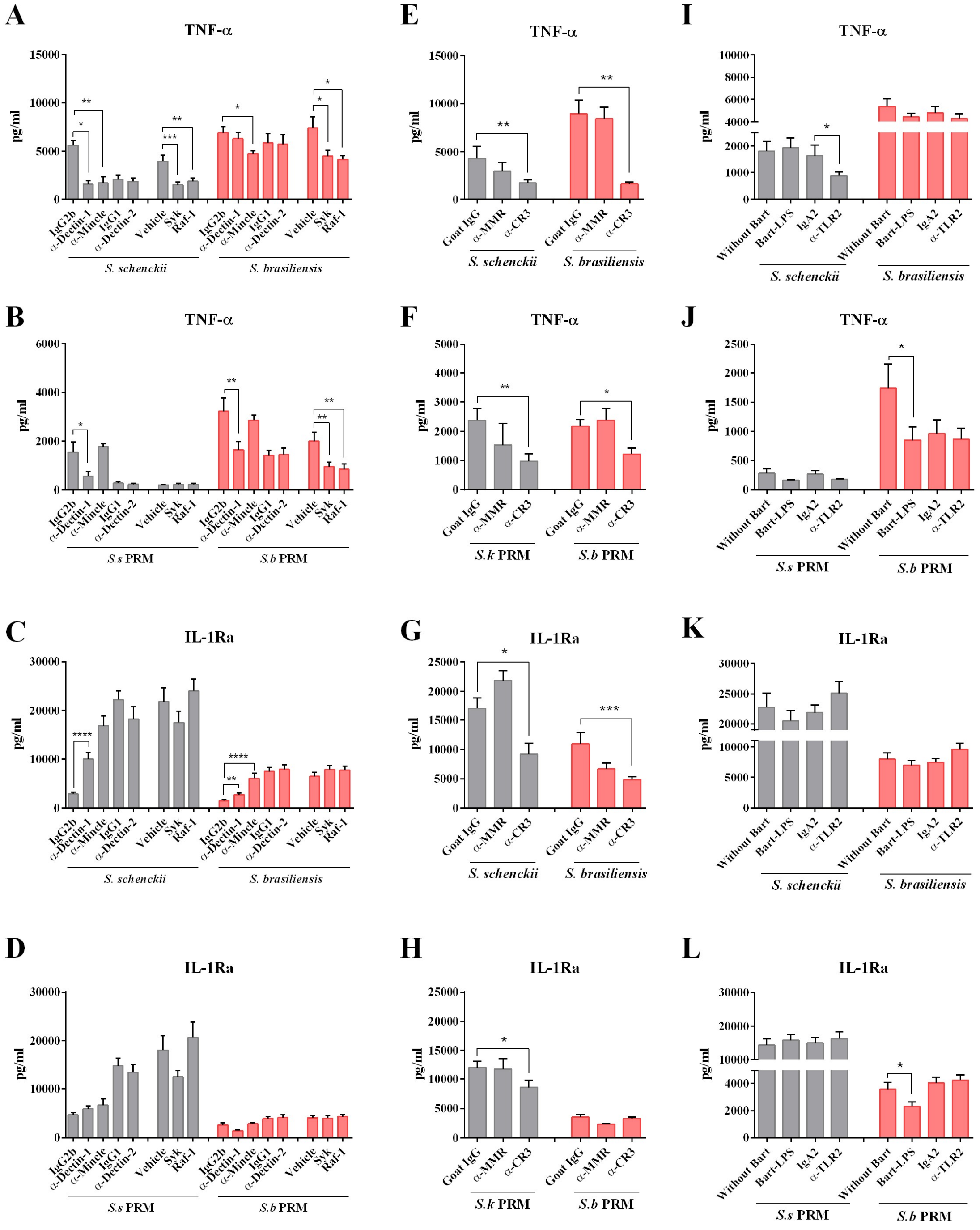


**Figure S2. C-type lectin, TLRs receptors and signalling pathways involved in the TNF-α and IL-1Ra production by *S. schenckii*, *S. brasiliensis* and their respective peptidorhamnomannans.** PBMCs were pre-incubated for 1 h with **A-D.** 10 µg/ml anti-dectin-1 (n=6), anti-dectin-2 (n=7) and anti-mincle (n=4) antibodies, or control isotypes IgG2b (n=6) and IgG1 (n=7). For assessing the signalling pathways, the cells were pre-incubated with 50 nM Syk (n=8), 1µM Raf-1 (n=8) and Vehicle (DMSO; n=8) as a control. **E-H.** anti-CR3 (n=8) and anti-MMR (n=4) antibodies, or isotype Goat IgG (n=8) as a control, **I-L.** anti-TLR2 (n=8) and 20 ng/ml Bartonella quintana LPS (n=8), or isotype IgA2 (n=8) and without Bart (RPMI only; n=8) as a control. After the blocking period, cells were stimulated with heat-killed *S. schenckii* and *S. brasiliensis* or *S.s* PRM and *S.b* PRM. After 24 hours of stimulation, supernatants were collected. TNF-α and IL-1Ra production were measured by ELISA. The data were expressed as mean ± SEM. Statistical analysis was performed by Wilcoxon test. *p < 0.05, **p < 0.01, ***p <0,001, ****p < 0.0001; differs from controls of isotype antibody or Vehicle.

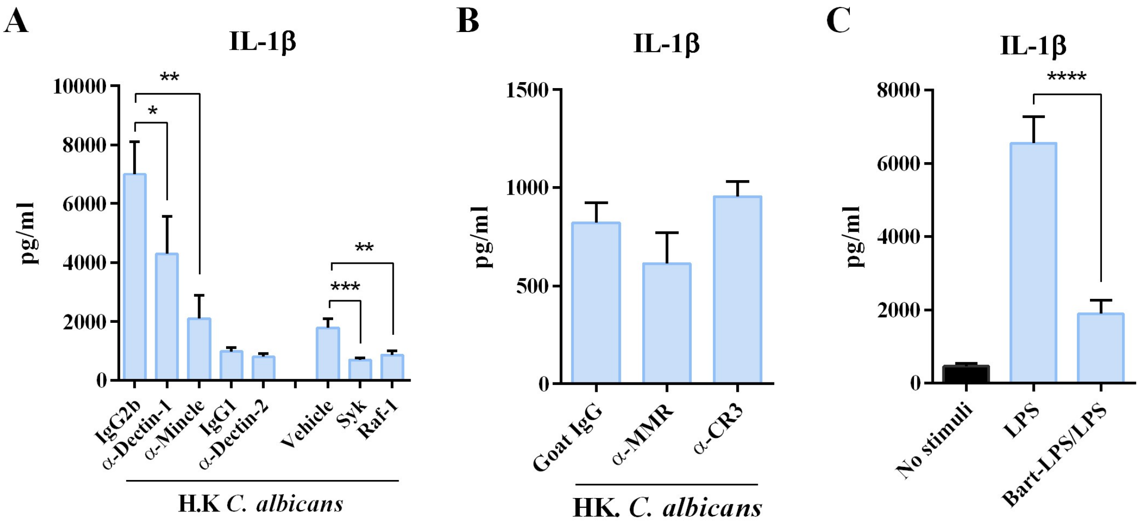


**Figure S3. Receptors and signalling pathways involved in the cytokine response by *C. albicans* and LPS as controls.** To block receptors, PBMCs were pre-incubated for 1 h with 10 µg/ml anti-dectin-1 (n=6), anti-dectin-2 (n=7) and (mincle n=4), anti-CR3 (n=8) and anti-MMR (n=4), anti-TLR2 (n=8) antibodies, or control isotypes IgG2b (n=6) and IgG1 (n=8), Goat IgG (n=8) IgA2 (n=8) and 20 ng/ml *Bartonella quintana* LPS (n=8). For signalling pathways the cells were pre-incubated with 50 nM Syk (n=8), 1µM Raf-1 (n=8) and Vehicle (DMSO; n=8) as a control. After the blocking period, cells were stimulated with heat-killed *C. albicans* or 10 ng/ml *Escherichia coli* LPS. After 24 hours of stimulation, supernatants were collected and IL-1β was measured by ELISA. Statistical analysis was performed by Mann-Whitney-U test. *p < 0.05, **p < 0.01, ***p < 0.001 ****p <0,0001; differs from controls isotype antibody or without Bart or vehicle.

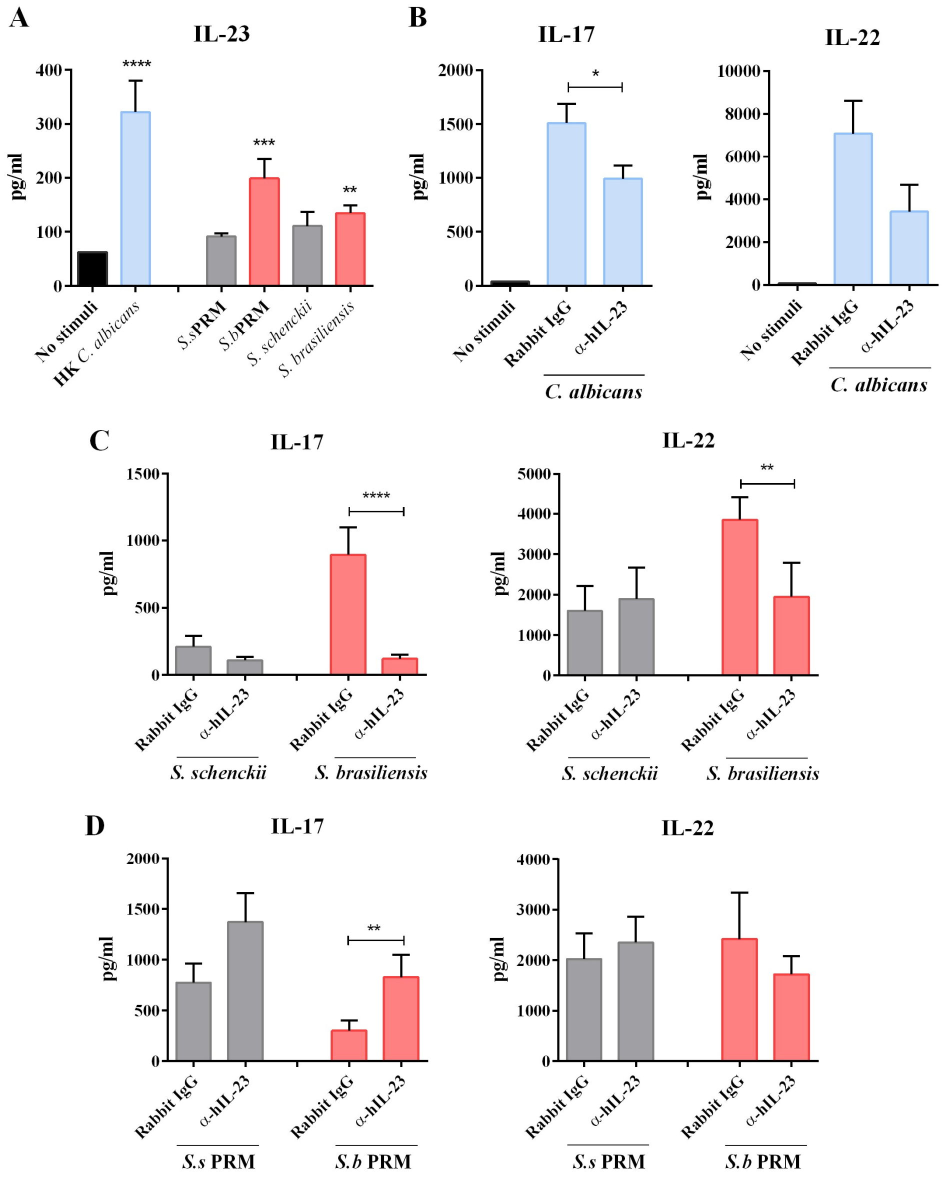


**Figure S4. IL-23 is important for the Th17 response of *Sporothrix brasiliensis* but not for peptidorhamnomannan. A.** PBMCs were stimulated with heat-killed *Candida albicans*, *S. schenckii* and *S. brasiliensis*. After 24 hours of stimulation, supernatants were collected and IL-23 (n=9) production were measured by ELISA. Statistical analysis was performed by Mann-Whitney-U test. $*p < 0.05$, $**p < 0.01$, $***p < 0{,}001$, $****p < 0.0001$; differs from no stimuli control. PBMCs were pre-incubated for 1 h with 10 µg/ml anti-hIL-23 or control isotypes Rabbit IgG (n=6). After the blocking period, cells were stimulated with $1x10^6$ cells/ml of heat-killed **B.** *C. albicans* as a control and **C.** *S. schenckii*, *S. brasiliensis* and **D.** 100 µg/ml of *S.s* PRM and *S.b* PRM. After 7 days of stimulation, supernatants were collected and IL-17 and IL-22 (n=6) production were measured by ELISA. Statistical analysis was performed by Mann-Whitney-U test. $*p < 0.05$, $**p < 0.01$, $****p < 0{,}0001$; differs from controls isotype antibody.

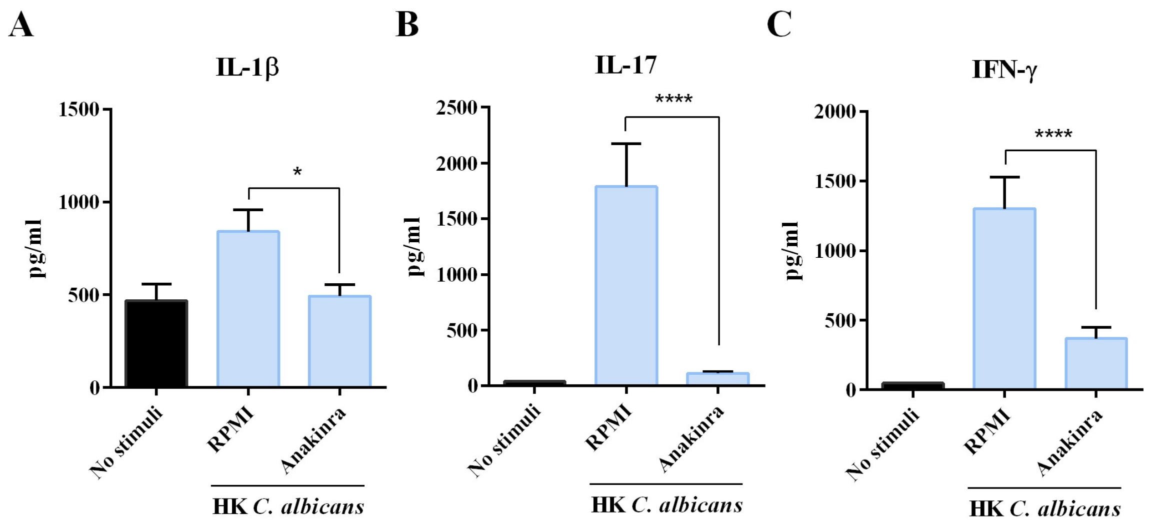


**Figure S5. The blockade of IL-1 directly impacts the Th1 and Th17 response of *C. albicans*.** PBMCs were pre-incubated for 1 h with and without 10 µg/ml Anakinra. After the blocking period, cells were stimulated with heat-killed *C. albicans*. Cells were incubated for 24 hours and 7 days at 37 ºC and 5% $CO_2$. After 24 hours of stimulation, supernatants were collected and **A.** IL-1β (n=8) production were measured by ELISA. After 7 days of stimulation, supernatants were collected and **B.** IL-17 and **C.** IFN-γ (n=9) production were measured by ELISA. Statistical analysis was performed by Mann-Whitney-U test. $*p < 0.05$, $**p < 0.01$, $***p < 0.001$ $****p < 0{,}0001$; differs from RPMI control (PBMCs without anakinra and treated with the respective stimulus).

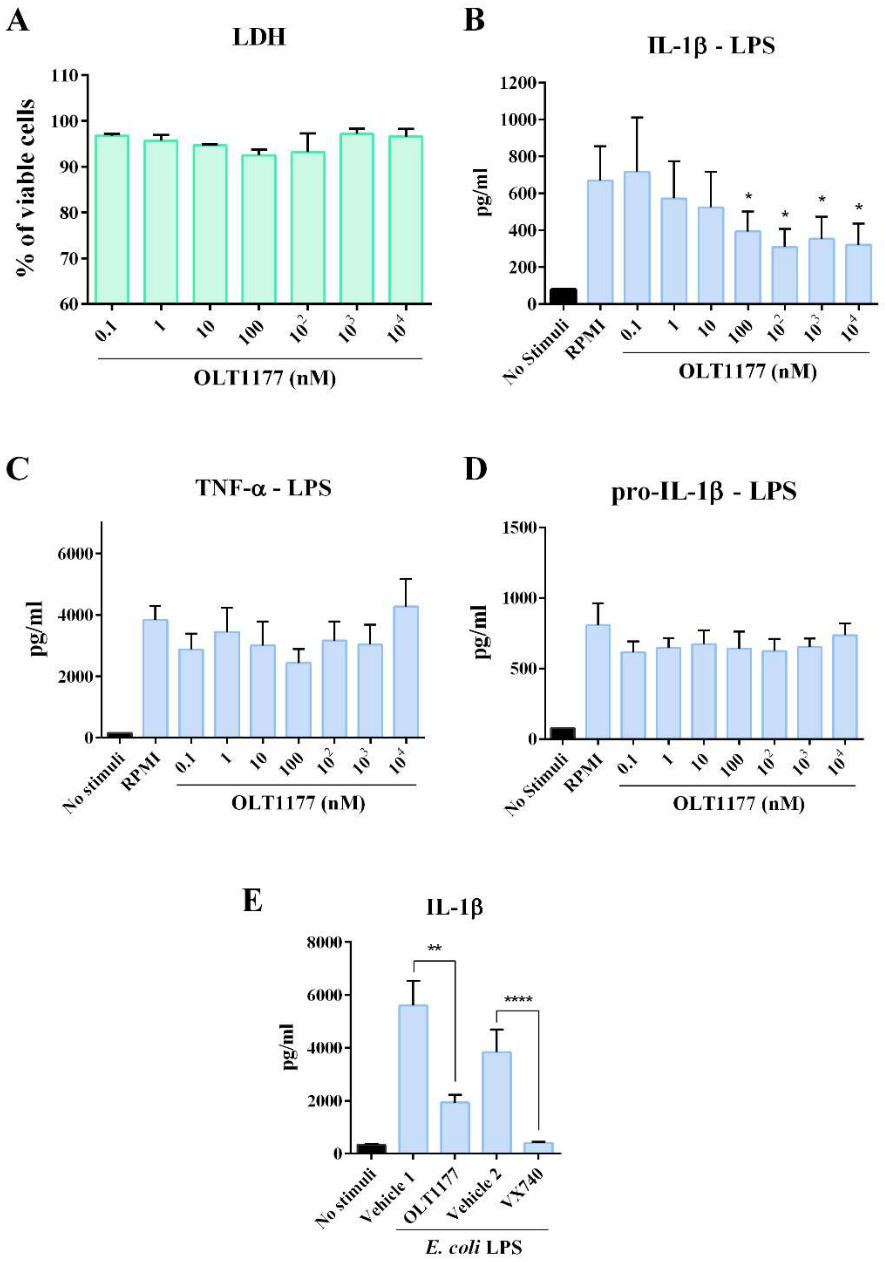


**Figure S6. OLT1177 is safe for PBMCs in adequate concentration to inhibit IL-β induction in LPS stimulated cells. A** PBMCs were pre-incubated for overnight with different concentrations of OLT1177 (0.1 – $10^4$ nM). After 24 h, LDH in the supernatants was measured by Cytotox96 non-Radioactive cytotoxicity assay. After the blocking period, cells were stimulated with 10 ng/ml *Escherichia coli* LPS. After 24 hours of stimulation, the supernatants were collected to measure **B.** IL-1β and **C.** TNF-α. The cells were lysate to measured **D.** pro-IL-1β by ELISA. **E.** PBMCs were treated overnight with OLT1177 or Pralnacasan (VX-740) and their respective vehicles as controls (Vehicle 1: PBS; Vehicle 2: DMSO). After 24 of stimulation with LPS, IL-b production was measured in the supernatants by ELISA. Statistical analysis was performed by Mann-Whitney-U test. ****p <0,0001; differs from the treatment with vehicle control.

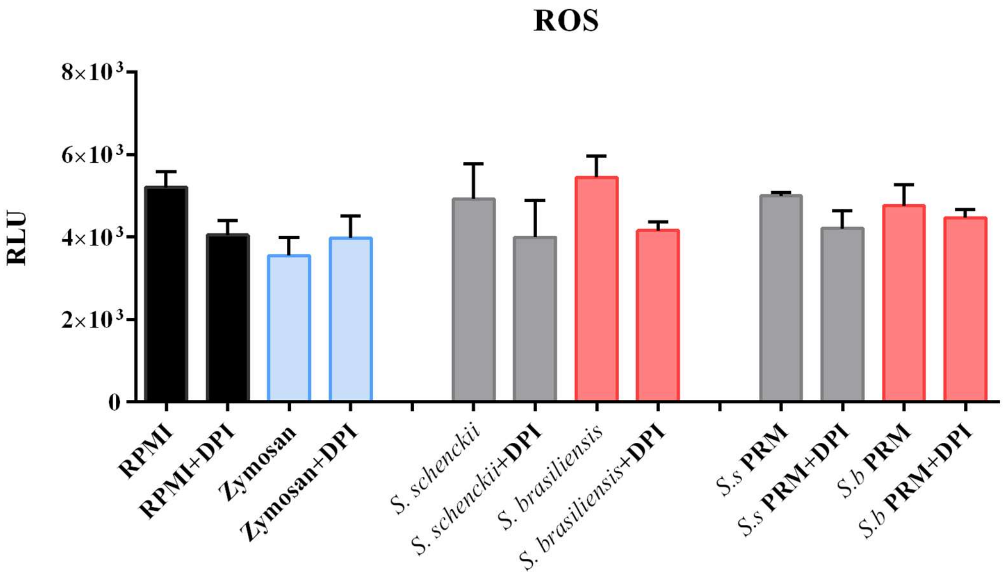


**Figure S7. Relative ROS production induced by *S. schenckii*, *S. brasiliensis* and their respective PRMs after 24h**. Total ROS production measured by area under each curve for the whole period of 1 hour. ROS production was measured by luminol-enhanced chemiluminescence assay. Cells were treated with 10 µM diphenyleneiodonium chloride (DPI) for 30 min and stimulated with Zymosan, *Sporothrix* yeast and PRMs as described above for 24 hours. The data were expressed as mean ± SEM. Statistical analysis was performed by Wilcoxon test. There was no statistical difference between the treated groups or in relation to the untreated control.
